# A two-step mechanism for the inactivation of microtubule organizing center function at the centrosome

**DOI:** 10.1101/446088

**Authors:** Jérémy Magescas, Jennifer C. Zonka, Jessica L. Feldman

## Abstract

**Summary:** During mitosis, the centrosome acts as a microtubule organizing center (MTOC), orchestrating microtubules into the mitotic spindle through its pericentriolar material (PCM). This activity is biphasic, cycling through assembly and disassembly during the cell cycle. Although hyperactive centrosomal MTOC activity is a hallmark of some cancers, little is known about how the centrosome is inactivated as an MTOC. Analysis of endogenous PCM proteins in *C. elegans* revealed that the PCM is composed of distinct protein territories that are removed from the centrosome at different rates and using different behaviors. Inhibition of PP2A phosphatases stabilized the PCM and perturbation of cortical pulling forces altered the timing and behavior by which proteins were removed from the centrosome. These data indicate that PCM disassembly is a two-step process, beginning with a phosphatase-dependent dissolution of PCM proteins followed by the ejection of ruptured PCM by cortical forces, ultimately inactivating MTOC function at the centrosome.

## Introduction

Numerous cell functions such as transport, migration, and division are achieved through the specific spatial organization of microtubules imparted by microtubule organizing centers (MTOCs). The best-studied MTOC is the centrosome, a membrane-less organelle composed of two barrel-shaped microtubule-based centrioles surrounded by a cloud of pericentriolar material (PCM). Microtubules at the centrosome are mainly nucleated and localized by complexes within the PCM, which generate a radial array of microtubules in dividing animal cells and some cell specialized types such as fibroblasts.

The PCM is a central hub for the regulation of numerous cellular processes including centriole duplication, ciliogenesis, cell cycle regulation, cell fate determination, and microtubule organization (Chichinadze, Lazarashvili, & Tkemaladze, 2012; Fry, Sampson, Shak, & Shackleton, 2017; Stubenvoll, Medley, Irwin, & Song, 2016). In *Drosophila* and human cell lines, PCM proteins are organized in cumulative layers to ultimately recruit microtubule nucleation and organization factors, such as the conserved microtubule nucleating γ-tubulin ring complex (γ-TuRC) (Fu & Glover, 2012; Lawo, Hasegan, Gupta, &Pelletier, 2012; Mennella, Agard, Huang, &Pelletier, 2014; Mennella et al., 2012). In *C. elegans*, the PCM is much simpler in composition, built from the interdependent recruitment of two scaffolding proteins, SPD-2/CEP192 and SPD-5 (Hamill, Severson, Carter, &Bowerman, 2002; Kemp, Kopish, Zipperlen, Ahringer, &O’Connell, 2004; Wueseke, Bunkenborg, Hein, Zinke, Viscardi, Woodruff, Oegema, Mann, Andersen, &Hyman, 2014b). Together with AIR-1/Aurora-A, SPD-2 and SPD-5 are required to localize γ-TuRC, which in *C. elegans* is composed of TBG-1/γ-tubulin, GIP-1/GCP3, GIP-2/GCP2 and MZT-1/MZT1 (Bobinnec, Fukuda, &Nishida, 2000; Hamill et al., 2002; Hannak et al., 2002; Kemp et al., 2004; Lin, Neuner, &Schiebel, 2015; Oakley, Paolillo, &Zheng, 2015; Sallee, Zonka, Skokan, Raftrey, &Feldman, 2018). γ-TuRC and AIR-1 together are required to build microtubules at the centrosome in the *C. elegans* zygote (Motegi, Velarde, Piano, &Sugimoto, 2006). Although the pathways required to build the PCM are largely known in *C. elegans*, the organization of proteins within the PCM has been unexplored.

The centrosome is not a static organelle; during each cell cycle, MTOC activity at the centrosome is massively increased to ultimately build the mitotic spindle (Dictenberg et al., 1998; Woodruff, Wueseke, &Hyman, 2014). This increase in centrosomal MTOC activity relies on the recruitment of PCM proteins to the centrosome, a process that is controlled by the concentration and availability of PCM proteins and their phosphorylation by mitotic kinases (e.g. CDK1, PLK1, and Aurora A) (Conduit et al., 2010; Conduit, Feng, et al., 2014a; Conduit, Richens, et al., 2014b; Decker et al., 2011; Novak, Wainman, Gartenmann, &Raff, 2016; Wueseke, Bunkenborg, Hein, Zinke, Viscardi, Woodruff, Oegema, Mann, Andersen, &Hyman, 2014a; Wueseke et al., 2016; Yang &Feldman, 2015). During mitotic exit, MTOC activity of the centrosome rapidly decreases, marked by the reduction of the PCM and microtubule association. Although the mechanisms controlling PCM disassembly have been relatively unexplored, inhibition of CDK activity can drive precocious PCM disassembly and inhibition of the PP2A phosphatase LET-92 perturbs SPD-5 removal from the centrosome, suggesting that phosphatase activity could be more generally required for the inactivation of MTOC function at the centrosome (Enos, Dressler, Gomes, Hyman, &Woodruff, 2018; Yang &Feldman, 2015). This cycle of centrosomal MTOC activity continues every cell cycle, but can also be naturally discontinued during cell differentiation when MTOC function is often reassigned to non-centrosomal sites (Sanchez &Feldman, 2016).

The inactivation of MTOC activity of the centrosome is likely critical in a number of cellular and developmental contexts. For example, asymmetric cell division is often associated with unequal PCM association at the mother vs. daughter centrosome and terminal differentiation of murine cardiomyocytes and keratinocytes has been linked to centrosome inactivation (Cheng, Tiyaboonchai, Yamashita, &Hunt, 2011; Conduit &Raff, 2010; Muroyama, Seldin, &Lechler, 2016; Zebrowski et al., 2015). In an extreme example, female gametes in a range of organisms completely eliminate centrosomes and this elimination can be a critical step in gametogenesis (Borrego-Pinto et al., 2016; Lu &Roy, 2014; Luksza, Queguigner, Verlhac, &Brunet, 2013; Mikeladze-Dvali et al., 2012; Pimenta-Marques et al., 2016). Moreover, hyperactive MTOC function at the centrosome has been linked to several types of epithelial cancers and invasive cell behavior, and is a hallmark of tumors (Godinho &Pellman, 2014; Lingle, Lutz, Ingle, Maihle, &Salisbury, 1998; Pihan, 2013; Pihan et al., 2001; Salisbury, Lingle, White, Cordes, &Barrett, 1999). Despite the clear importance of properly regulating MTOC activity, little is known about the mechanisms that inactivate MTOC function at the centrosome, either what initiates the removal of PCM and microtubules during the cell cycle or what keeps them off the centrosome in differentiated cells.

To better understand how MTOC activity is regulated at the centrosome, here we investigate the localization and dynamics of endogenously tagged PCM proteins in the *C. elegans* embryo. We find that *C. elegans* PCM is composed of layered spheres of proteins, with SPD-5 and γ-TuRC occupying distinct regions from known binding partners SPD-2 and MZT-1, respectively. Live imaging of SPD-2, SPD-5, and γ-TuRC components at the end of mitosis revealed two phases of disassembly, beginning with the gradual dissolution of PCM proteins, followed by the rupture of the remaining PCM into small microtubule associated packets. Using pharmacological or genetic perturbations, we found a role for PP2A phosphatases in the initial dissolution of PCM proteins and for cortical pulling forces in the clearance of the remaining PCM from the centrosome. Delay in PCM removal impacted subsequent centriole separation and PCM maturation in the next cell cycle. These data indicate that the inactivation of MTOC function at the centrosome involves a regulated two-step process of PCM disassembly, the timing of which is critical to the developing embryo.

## Results

### *C. elegans* PCM is organized into an inner and outer sphere

In order to better understand how PCM proteins behave during disassembly, we first characterized the spatial organization of the PCM during mitosis in the ABp cell of the 4-cell *C. elegans* embryo. ABp has relatively large centrosomes oriented during cell division along the left-right axis of the embryo, with one of the centrosomes positioned very close to the coverslip with an end-on orientation (Figure 1A). We analyzed the localization of endogenously-tagged PCM proteins immediately after nuclear envelope breakdown (NEBD) in the ABp cell (Figure 1A) (Dickinson, Pani, Heppert, Higgins, &Goldstein, 2015; Dickinson, Ward, Reiner, &Goldstein, 2013). At this time, the centrosome still functions as an MTOC, actively growing and organizing microtubules (Figure 1A).

**Figure 1.**
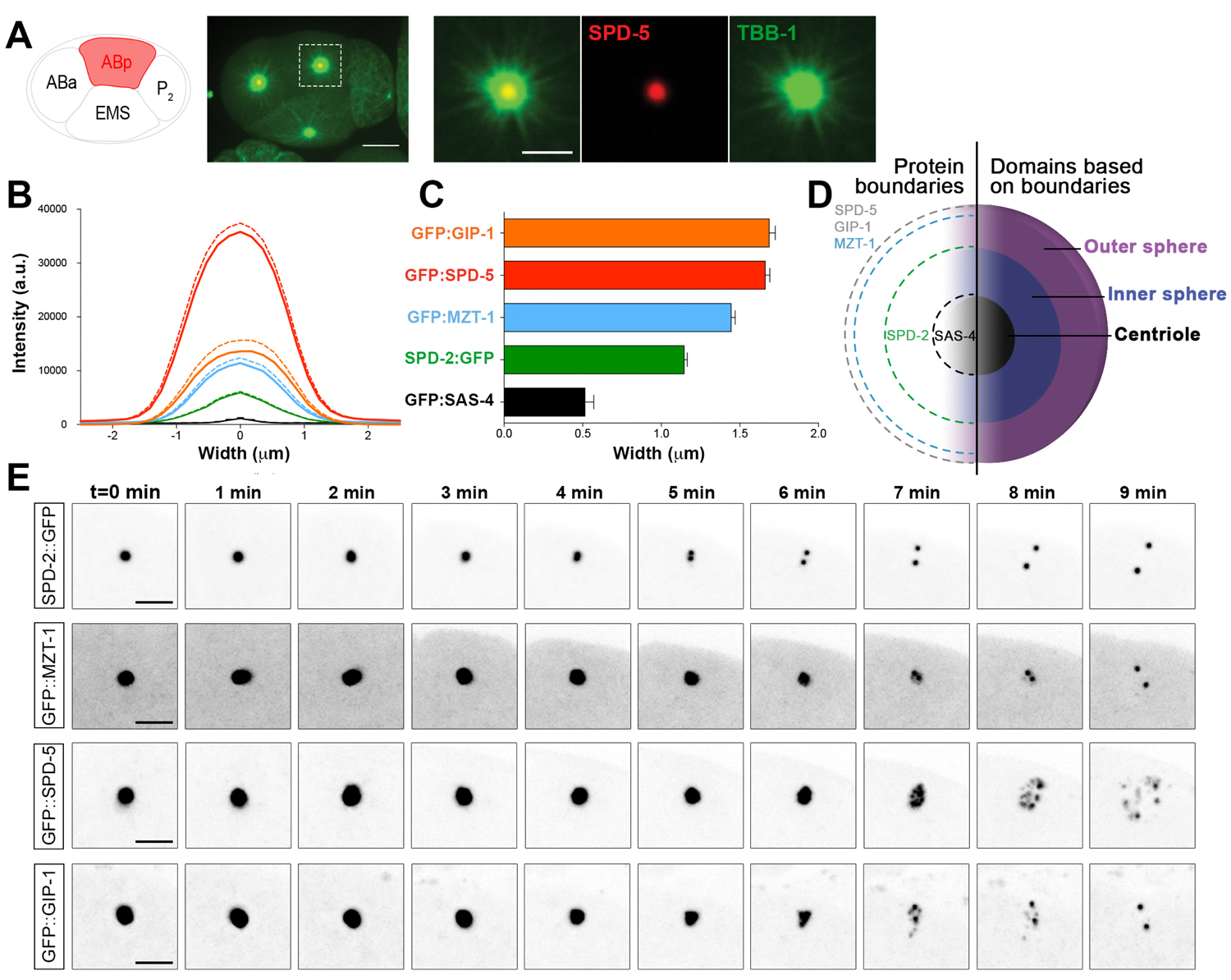
*C. elegans* PCM is organized into two spheres that disassemble using different behaviors, see also Figure 1 - figure supplement 1, Videos 1 to 4. A) Left: Cartoon representing the *C. elegans* 4-cell stage embryo with ABp in red. Right: 7.5μm z-projection from a live *pie-1p*::GFP::TBB-1/β-tubulin (green); tagRFP::SPD-5 (red) expressing embryo showing cell division in ABa and ABp. Note that these cells have a synchronized cell division and start dividing earlier than EMS or P2. Insets: Enlargement of ABp centrosome showing microtubules (green) organized around the centrosome (SPD-5, red). Scale bar, 5 μm. B) Average pixel intensity profile across the ABp centrosome at NEBD: GFP::GIP-1 (orange, n=18), GFP::SPD-5 (red, n= 18), GFP::MZT-1 (blue, n= 21), SPD-2::GFP (green, n= 21), GFP::SAS-4 (black, n=19). Bold line represents the mean, dotted lines represent standard error of the mean (s.e.m.). C) Average width of pixel intensity profile for each protein in B. GIP-1: 1.69±0.04|jm, n=18; SPD-5: 1.66±0.03|μm, n= 18; MZT-1: 1.44±0.03μm, n= 21; SPD-2: 1. 15±0.02μm, n= 21; SAS-4: 0.51±0.06μm, n= 19. D) Cartoon representing the organization of the centrosome based on the boundary of SAS-4 (black, “centriole”), SPD-2 (dark blue, “inner sphere”), and SPD-5 (purple, “outer sphere”. E) Time-lapse analysis of the disassembly of each protein analyzed in B, C and D starting at NEBD (t=0 min) and imaged every minute for 9 minutes. Scale bar, 10μm.

We assessed the localization of the centriole component SAS-4, the PCM proteins SPD-2 and SPD-5, and the γ-TuRC components GIP-1 and MZT-1 (Figure 1B, Figure 1 - supplement 1). As expected, the centrioles sit at the center of the centrosome (Figure1B, C) surrounded by SPD-2, SPD-5 and γ-TuRC. Interestingly, SPD-2 and SPD-5 displayed two distinct boundaries, with both SPD-2/CEP192 and SPD-5 localizing to a more internally restricted “inner sphere” (Figure 1B, C) and SPD-5 extending further into an “outer sphere” (Figure 1B, C). Similar to SPD-5, GIP-1 localization extended into the outer sphere (Figure 1B, C), however surprisingly, the γ-TuRC component MZT-1 showed an intermediary localization, extending to a region between the inner and outer sphere (Figure 1B, C). Based on these observations, we conclude that the PCM has a layered structure with an inner sphere delimited by SPD-2 (Figure 1D) that also localizes SPD-5 and γ-TuRC components, and an outer sphere delimited by the SPD-5 and GIP-1 (Figure 1D). This organization follows the general pattern of the predicted orthologs in *Drosophila* and human cells (Fu &Glover, 2012; Lawo et al., 2012; Mennella et al., 2012; 2014), but is somewhat surprising as SPD-5 and GIP-1 are found in a region lacking known binding partners SPD-2 and MZT-1, respectively.

### PCM proteins disassemble with different behaviors

Based on their distinct localization within the PCM, we hypothesized that different PCM proteins would disassemble with different kinetics and behaviors. To test this hypothesis, we examined the dynamics of disassembly of each of the endogenously-tagged PCM proteins described above by live-imaging in the ABp cell beginning at NEBD (Figure 1E). SPD-2 (Video 1) and MZT-1 (Video 2) displayed similar disassembly behaviors, leaving the centrosome by gradual dissolution over time. In contrast, SPD-5 (Video 3) and GIP-1 (Video 4) initially showed a gradual pattern of disassembly, however the structure containing these proteins then appeared to rupture and fragment into “packets” that were distinct from the centrioles. These sub-PCM packets localized SPD-5, GIP-1/GCP3 (Figure 2A, *early packets*), and microtubules (Figure 2B, *early packets*), but neither SPD-2 nor MZT-1 (Figure 2D, see below). Intriguingly, packets appeared to retain MTOC potential as EBP-2/EB1 comets, a marker of growing microtubule plus ends, dynamically moved from the SPD-5/GIP-1 foci (Figure 2C). The packets appeared to be further disassembled in the cytoplasm following their removal from the PCM, with GIP-1 and microtubules first losing their association, followed by SPD-5 (Figure 2A,B, *late packets*, Figure 2E).

**Figure 2.**
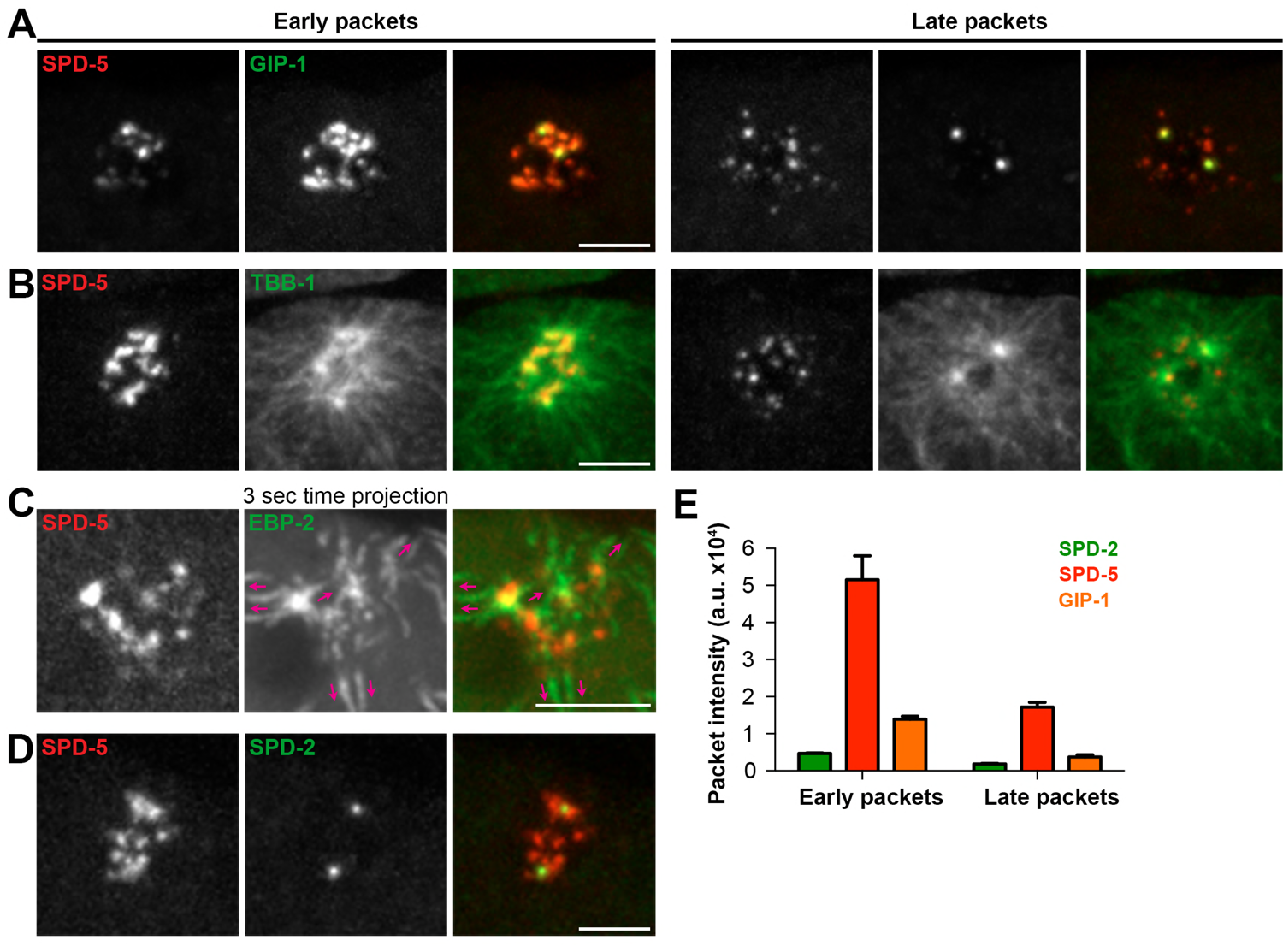
The PCM fragments into SPD-5 and GIP-1 containing packets that localize dynamic microtubules. A-B) Analysis of colocalization of SPD-5 packets (red) with GIP-1 (A, green), or microtubules (B, TBA-1/α-tubulin, green) in early packets (left panels) or late packets (right panels). C) Three second time projection of EBP-2 (green) showing that packets (SPD-5, red) associate with dynamic microtubules. Magenta arrows represent the orientation of EBP-2 movement. Scale bar, 10μm. D) Colocalization of SPD-5 packets (red) with SPD-2 (green). Note that SPD-2 does not localize to the packets. E) Average pixel intensity of SPD-2 (green, n=8), SPD-5 (red, n=11), and GIP-1 (orange, n=8) in early and late packets. ‘a.u.’ = arbitrary units. Graph represent mean ± s.e.m.

To gain a better sense of the timing of the disassembly of the different PCM proteins, we imaged each protein in combination with SPD-5. SPD-2 (Figure 3A) and MZT-1 (Figure 3B) showed a gradual decrease in intensity, beginning at 2 (2.20±0.13 min, n=10) or 3 minutes (3.00±0.27 min, n=8) post-NEBD, respectively, several minutes before the decrease in either SPD-5 or GIP-1 (Figure 3D, E). Consistent with this trend, we found that the PCM volume of SPD-2 and MZT-1 gradually decreased beginning 3 minutes post-NEBD (SPD-2: 3.00±0.21 min, n=10; MZT-1: 2.88±0.23 min, n=8). As expected from our observation of the individual localization behaviors, both SPD-5 and GIP-1 co-localized during the process of disassembly (Figure 3C, Video 5). We observed the same trend in both the total intensity and PCM volume of SPD-5 and GIP-1 (Figure 2D, E); both proteins rapidly decreased in intensity following their peak at 3 minutes post-NEBD (SPD-5: 3.00±0.14 min, n=11; GIP-1: 3.18±0.12 min, n=11), and their volume dramatically and precipitously reduced at the PCM beginning 6 minutes post-NEBD (SPD-5: 5.91±0.17 min, n=11; GIP-1: 6.00±0.19 min, n=11), reflecting packet formation. Together, these data indicate that the PCM disassembles in two distinct steps: a dissolution step that is characterized by the decrease in intensity of PCM proteins that starts with the removal of SPD-2 and MZT-1; and a rupture/packet formation step where the deformation and subsequent rupture of the PCM leads to further disassembly into individual packets.

**Figure 3.**
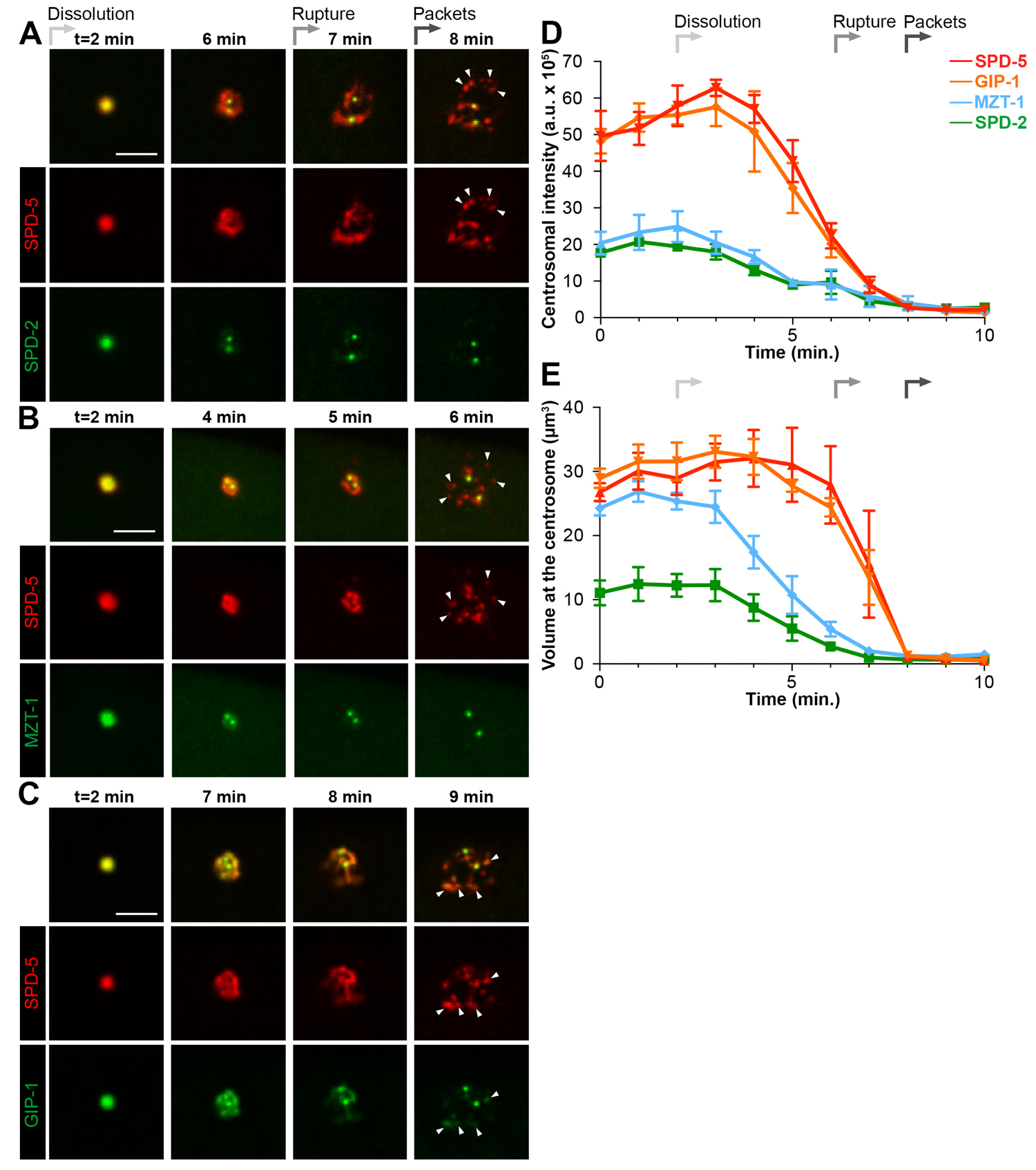
Loss of SPD-2 and MZT-1 precedes rupture and packet formation, see also Video. 5 A-C) Comparison of tagRFP::SPD-5 (red) to SPD-2::GFP (A, green), GFP::MZT-1 (B, green), GFP::GIP-1 (C, green) disassembly. ‘Dissolution’ (light grey arrow) begins as SPD-2 (t=2 min. post-NEBD) and then MZT (t=3 min post-NEBD) are removed from the centrosome. ‘Rupture’ (medium grey arrow) is indicated by holes appearing in the matrix of SPD-5 and GIP-1 surrounding the centrioles, followed by the appearance of individual ‘packets’ (dark grey arrow) of SPD-5 and GIP-1. Scale bar, 10μm. D-E) Average pixel intensity (D) and volume (E) at the centrosome of PCM proteins during disassembly starting at NEBD (t=0 min): tagRFP::SPD-5 (red, n= 11), GFP::GIP-1 (orange, n=9), GFP::MZT-1 (sky blue, n= 10), SPD-2::GFP (green, n= 8). ‘a.u.’ = arbitrary units. Graph lines indicate mean ± s.e.m.

### Cortical forces mediate the disassembly of the PCM and more specifically SPD-5

The formation of packets that appear to be pulled away from the centrioles suggests that mechanical forces underlie this aspect of PCM disassembly. Forces can be exerted on the PCM by a conserved cortically anchored complex of LIN-5/NuMA, LGN/GPR-1/2, and (GOA-1/GPA-16)/Gαi, which localizes dynein-dynactin that can pull on the astral microtubules extending from the PCM (Kotak &Gönczy, 2013). Given that greater cortical forces exist in the posterior of the one-cell *C. elegans* embryo, it has been hypothesized that these forces could be responsible for the asynchrony observed in the disassembly of the anterior vs. the posterior centrosome (Grill, Gönczy, Stelzer, &Hyman, 2001). Moreover, a recent study implicated the GPR-1/2/LIN-5/DHC-1 complex in SPD-5 disassembly from the PCM (Enos et al., 2018).

To assess the involvement of cortical forces in the general disassembly of the PCM and specifically in rupture and packet formation, we used RNAi to either decrease (*gpr-1/2*(RNAi)) or increase (*csnk-1*(RNAi)) cortical forces. In control embryos treated with lacZ RNAi, SPD-5 ruptured starting 6 min post-NEBD (5.91±0.16 min, n=11) and formed packets at 8 min post-NEBD (7.73±0.14 min, n=11; Figure 4A). In contrast, we did not observe SPD-5 rupture or packet formation in *gpr-1/2*(RNAi) treated embryos (Figure 4A). Instead, SPD-5, like SPD-2, was removed from the centrosome by gradual dissolution (Figure 4B). In *csnk-1*(RNAi) treated embryos, we observed slightly earlier SPD-5 rupture (5.4+0.2 min, n=7) and packet formation (7.14±0.14 min, n=7; Figure 4A). In contrast to SPD-5, SPD-2 disassembly appeared unaffected following depletion of either *gpr-1/2* or *csnk-1* by RNAi (Figure 4C). Interestingly, both SPD-5 intensity and volume were increased or decreased by either *gpr-1/2* or *csnk-1* depletion (Figure 4A, Figure 4 – figure supplemental 1). Together, these results suggest that cortical forces generate the mechanical forces necessary for rupture and packet formation, allowing for the removal of the outer sphere protein SPD-5 but not the exclusively inner sphere protein SPD-2.

**Figure 4.**
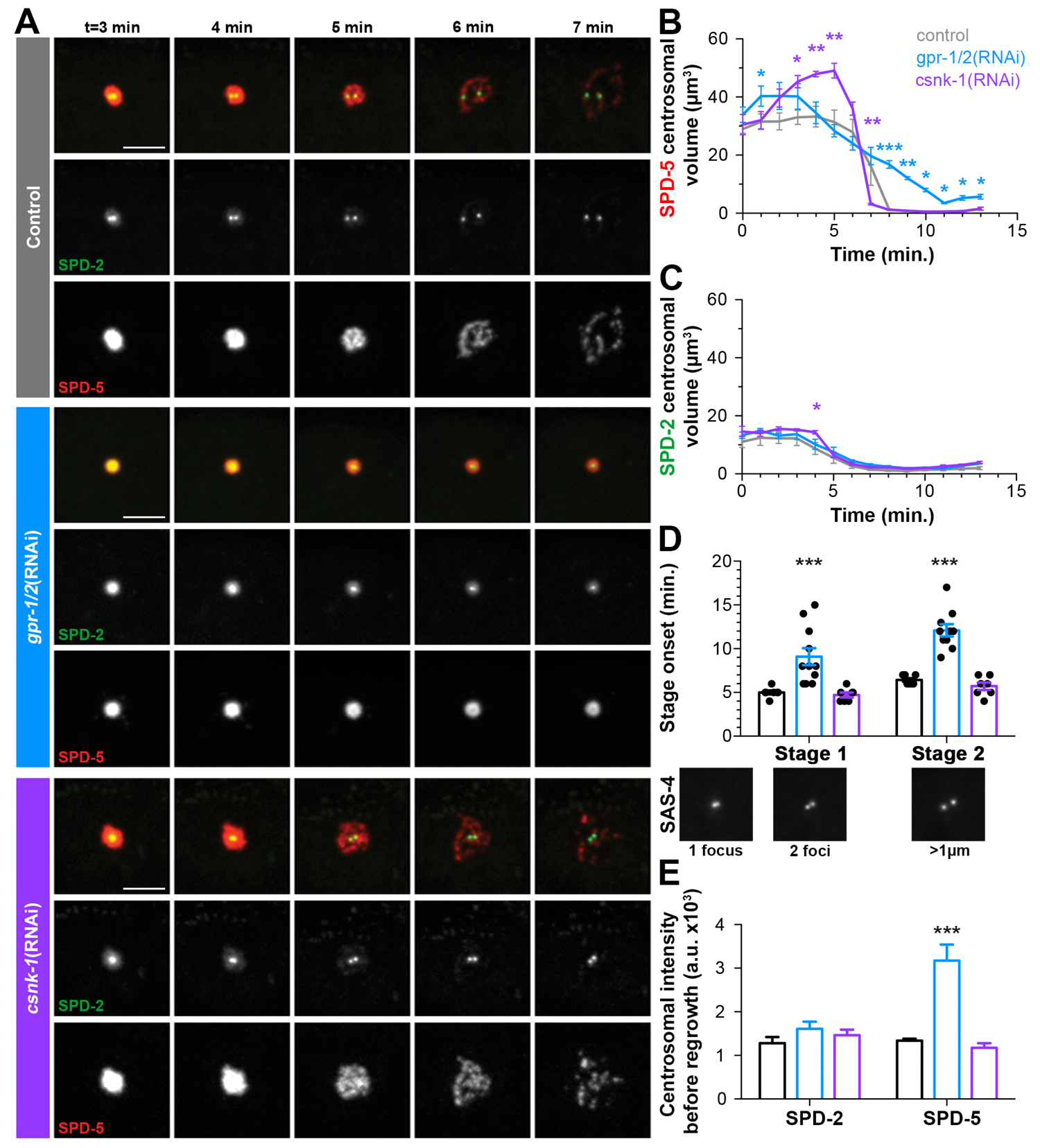
Cortical forces rupture the PCM into packets, see also Figure 4 – figure supplement 1 and supplement 2. A) Time-lapse analysis starting at NEBD (t=0 min) of the disassembly of endogenous tagRFP::SPD-5 (red) and SPD-2::GFP (green) treated with lacZ(RNAi) (control, top panels, grey (A-E)), *gpr-1/2*(RNAi) (middle panels, blue (A-E)), or *csnk-1(RNAi)* (bottom panels, purple (A-E)). Scale bar, 10μm. B-C) Average volume at the centrosome of SPD-5 (B) or SPD-2 (C) during disassembly starting at NEBD (t=0 min). D) Average onset time for centriole separation starting at NEBD (t=0 min). Stage 1: Centrioles are apparent as a single focus and then double foci of GFP::SAS-4. Stage 2: Centrioles appear >1μm apart. control, Stage 1: 5.00±0.218 min; control, stage 2: 6.429±0.202 min, n=8; *gpr-1/2*(RNAi), Stage 1: 9.091±0.977 min, *gpr-1/2*(RNAi), Stage 2: 12.100±0.706 min, n=11; *csnk-1*(RNAi), Stage 1: 4.714±0.286 min, *csnk-1*(RNAi), Stage 2: 5.714±0.421 min, n=7. E) Average intensity of SPD-2 or SPD-5 remaining at the centrosome before regrowth in the next cell cycle. SPD-2(control): 1281 ±139, SPD-5(control): 1337±47, n=8; SPD-2(*gpr-1/2*(RNAi)): 1610±166, SPD-5(*gpr-1/2*(RNAi)): 3173±369, n=11; SPD-2(csnk-1 (RNAi))/ 1467±122, SPD-5(*csnk-1*(RNAi)): 1172±110, n=7. Asterisks indicate comparison between indicated perturbation and control: *p-value < 0.01, ** p-value < 0.001, *** p-value < 0.0001. ‘a.u.’ = arbitrary units. Graphs indicate mean ± s.e.m.

Cortical forces could be present and constant throughout mitosis or instead intensify at the time of disassembly as is the case in the zygote, providing forces only when necessary (Gönczy &Rose, 2005; Rose &Gönczy, 2014). To distinguish between these possibilities, we tracked the localization of LIN-5, DNC-1/dynactin, DHC-1/dynein heavy chain, and microtubules during different stages of mitosis. We saw no change in the gross cortical distribution or intensity of LIN-5 (Figure 4 - figure supplement 2A) or DNC-1 (Figure 4 – figure supplement 2B) post-NEBD. Similarly, DHC-1 cortical localization appeared consistent over time (Figure 4 - figure supplement 2C), although we saw an ephemeral redistribution of DHC-1 coincident with rupture (Figure 4 - figure supplement 2D, 4 min.). Strikingly, astral microtubules showed a network reorganization post-NEBD, growing progressively longer and contacting the cell cortex, sometimes wrapping around the membrane (Figure 4 - figure supplement 2E). This pattern of localization suggests that although cortical complexes are present throughout the cell cycle, they may only make productive contact with astral microtubules at a particular time period to allow for outer sphere disassembly.

The rapid rounds of PCM assembly and disassembly during the early embryonic divisions suggest that efficient and robust PCM disassembly might be critical for subsequent carefully timed events such as centriole separation and the assembly of new PCM in the next cell cycle (Cabral, Sans, Cowan, &Dammermann, 2013). We tested whether force dependent PCM removal corresponds to centriolar separation by tracking SAS-4::GFP during disassembly (Figure 4D). In control embryos, the centriolar pair appeared as a single SAS-4 focus up to 5 minutes post-NEBD (Figure 4D). Two closely apposed SAS-4 foci became apparent beginning at 5 min post-NEBD (Stage 1, Figure 4D), which quickly separated by greater than 1μm beginning about 1 minute later (Stage 2, Figure 4D). We saw a significant delay in the onsets of both Stage 1 and Stage 2 in *gpr-1/2*(RNAi) treated embryos, but no significant change in csnk-1(RNAi) treated embryos (Figure 4D). These results suggest that cortical forces facilitate centriole separation either through direct force transmission or indirectly through their role in PCM removal. That *csnk-1* RNAi had no effect on the timing of centriole separation suggests that a force-independent licensing event is necessary to initiate separation (Cabral et al., 2013; Tsou &Stearns, 2006), but that centrioles are subsequently held together by PCM. In addition to defects in centriole separation, we observed that *gpr-1/2*(RNAi) treated embryos had defects in effectively clearing SPD-5, but not SPD-2, from the PCM prior to the subsequent round of PCM accumulation in the next cell cycle (Figure 4B, E). Consistent with these defects, the timing of subsequent SPD-5 accumulation was significantly delayed as compared to control embryos (Figure 4 - supplement 1C). Together, these results underscore the importance of the timely removal of PCM to the developing embryo.

### PP2A phosphatases are required for PCM dissolution

As the growth of the PCM is highly dependent on phosphorylation and CDK inhibition causes precocious removal of PCM proteins (Woodruff et al., 2014; Yang &Feldman, 2015), we hypothesized that the dissolution of the PCM that precedes rupture and packet formation requires phosphatase activity. To test this hypothesis, we treated cycling embryonic cells at anaphase with either a broad-spectrum serine/threonine phosphatase inhibitor (okadaic acid) or a PP2A inhibitor (rubratoxin A, Figure 5A). We observed a stabilization of the PCM in both okadaic acid and rubratoxin A treated embryos compared to control embryos treated with DMSO. Notably, treatment with either drug led to depolymerization of the microtubules, perhaps due to the hyperactivation of the depolymerizing kinesin KLP-7 during PP2A inactivation (Schlaitz et al., 2007). Consistent with these pharmacological inhibition results, a recent study implicated the PP2A subunit LET-92 in SPD-5 disassembly (Enos et al., 2018).

**Figure 5.**
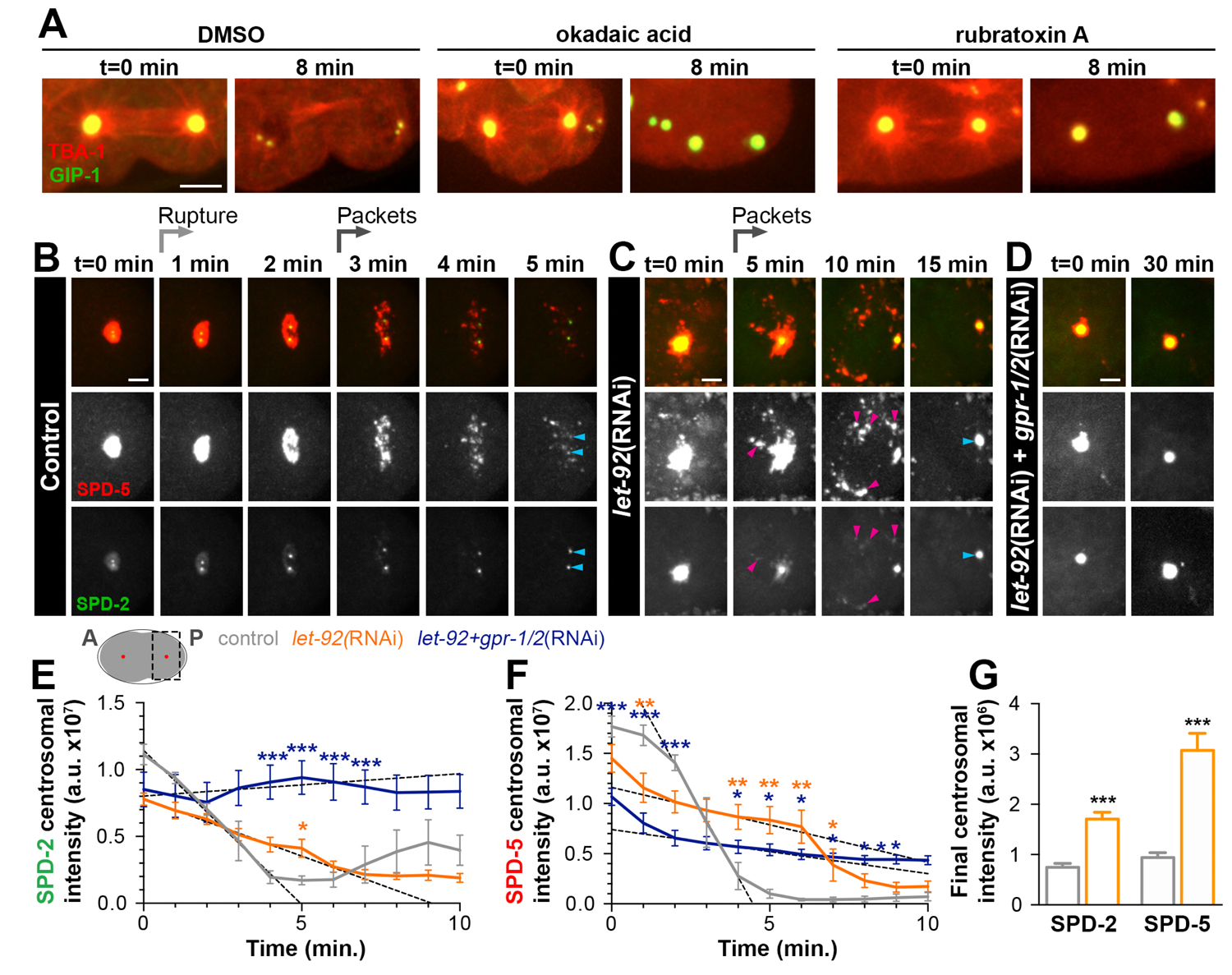
PP2A phosphatases regulate PCM disassembly. A) Time-lapse analysis of embryos expressing *pie-1p*::mCherry::TBA-1/α-tubulin (red) and endogenous GFP::GIP-1 (green) and treated at anaphase (t=0 min) with DMSO (left panels), 30 μM okadaic acid (middle panels), or 60 μM rubratoxin A (right panels). Scale bar, 10μm. B-D) Time-lapse analysis of the disassembly of endogenous tagRFP::SPD-5 (red); SPD-2::GFP (green) starting from cytokinetic furrow ingression (t=0 min) in the one cell embryo as represented on the cartoon below. Timing of rupture (light gray arrow) and packet formation (dark gray arrow) are indicated. Images show posterior (P) embryonic region (black dotted box in cartoon) containing the posterior centrosome (red dot in cartoon). Embryos are treated with lacZ(RNAi) (control, B), *let-92*(RNAi) (C), or *let-92*(RNAi) + *gpr-1/2*(RNAi) (D). Note the appearance of SPD-2 in packets (C, magenta arrowheads) following *let-92* RNAi treatment. Scale bars, 10μm. E-F) SPD-2 (E) or SPD-5 (F) intensity at the centrosome during disassembly starting from cytokinetic furrow ingression (t=0 min) in embryos treated with lacZ(RNAi) (control, grey, n=8), *let-92*(RNAi) (orange, n=8), or *let-92*+*gpr-1/2*(RNAi) (navy, n=8). SPD-2 disassembly slope (E, 0 to 4min, black dotted lines): control (slope=-2.31e+^6^, r^2^=0.97), *let-92*(RNAi) (slope=-8.60e+^5^, r^2^=0.94) and *let-92*+*gpr-1/2*(RNAi) (slope=1.67e+^5^, r^2^=0.86). SPD-5 disassembly slope (F, 2 to 4min, black dotted lines): control (slope=-5.65e+^6^, r^2^=0.95), let-92(RNAi) (slope=-7.46e+^5^, r^2^=0.92) and *let-92*+*gpr-1/2*(RNAi) (slope=-4.40e+^5^, r^2^=0.91). Slopes are significant different from each other using a t-test, p-value < 0.0001. G) Average centrosomal pixel intensity at the end of disassembly in control (t = 5.2 ± 0.2 min, grey, n=15) and in *let-92*(RNAi) (t = 12.1 ± 0.9 min, orange, n=13) treated embryos. Note that we accounted for centriole duplication defects following *let-92* depletion by comparing the average intensity of each individual centriole/centrosome in control embryos (see two SPD-2 foci representing two individual centrioles/centrosomes, light blue arrowheads at t = 5’ in Figure 5B) to intensity of the single centrosome in *let-92* depleted embryos (single SPD-2 focus, light blue arrowhead at t = 15’ in Figure 5C; see Material and Methods). Asterisks indicate comparison between indicated perturbation and control: *p-value < 0.01, ** p-value < 0.001, *** p-value < 0.0001. ‘a.u.’ = arbitrary units. Graphs indicate mean ± s.e.m.

To assess the function of LET-92 on PCM disassembly in general and more specifically on dissolution and packet formation, we treated SPD-2::GFP; tagRFP::SPD-5 expressing embryos with *let-92*(RNAi). As previously reported, *let-92* inhibition caused severe defects in cell division, necessitating analysis in the one-cell zygote rather than 4-cell embryo (Song, Liu, Anderson, Jahng, &O’Connell, 2011). We monitored PCM disassembly in the one-cell zygote beginning when the membrane invagination that occurs during cytokinetic furrow formation was visible. At this stage in control embryos, PCM disassembly occurs in a similar manner to ABp cells, with SPD-2 dissolution preceding SPD-5 rupture and packet formation (Figure 5B).

*let-92* depletion impaired the disassembly of SPD-2 and SPD-5 from the centrosome in three distinct ways (Figure 5C). First, SPD-5 was still partially cleared from the centrosome into packets, which persisted significantly longer in the cytoplasm as compared to control (Figure 5C). Interestingly, unlike in control embryos where SPD-2 was cleared from the centrosome by gradual dissolution, SPD-2 ruptured and frequently appeared in packets following *let-92* depletion (Figure 5C). Second, the rate and time of SPD-2 and SPD-5 disassembly were significantly slower in *let-92* depleted embryos than in control, as indicated by tracking the total centrosomal SPD-2 and SPD-5 over time (Figure 5E, F). Centriole duplication fails following *let-92* depletion such that each centrosome at this stage contains only one rather than two centrioles (Song et al., 2011). Thus, total centrosome intensity measurements underestimate differences between control and *let-92* depletion conditions because centriole number defects alter the underlying amounts of centriole-localized SPD-2 or SPD-5. Finally, we found that although much of the SPD-2 and SPD-5 appeared to be cleared from the PCM into packets, *let-92* depletion inhibited the complete removal of either protein from the centrosome (Figure 5C, G).

The partial removal of SPD-2 and SPD-5 in packets suggested that *let-92* depletion affected mainly dissolution, and that much, but not all, of the remaining PCM was cleared by cortical forces. To test this model, we inhibited *let*-92 together with *gpr-1/2* and observed a strong stabilization of both SPD-2 (Figure 5D, E) and SPD-5 (Figure 5D, F) at the PCM without rupture or packet formation. These data indicate that cortical forces are necessary but not sufficient to remove both SPD-2 and SPD-5 from the centrosome. Together, these results indicate that PP2A phosphatases control the dissolution of SPD-2 and SPD-5, and that both PP2A and cortical forces are required for the efficient and timely removal of the PCM from the centrosome.

## Discussion

We found that the *C. elegans* centrosome is organized into an inner and an outer sphere of PCM, which disassemble via a two-step mechanism. This organization appears to be generally conserved between direct and functional orthologs in *C. elegans*, Drosophila, and human PCM, suggesting evolutionary pressure to create specific functional PCM domains and that the mechanisms of disassembly described here might be generally conserved. The existence of SPD-5 and GIP-1 in a region lacking SPD-2 and MZT-1 suggests that these proteins have the ability to form a matrix in the absence of their known binding partners. SPD-5 can form a matrix in vitro and perhaps its self-association drives outer sphere assembly (Woodruff et al., 2015). Similarly, experiments from *S. pombe* suggests that MZT1 drives the assembly of the γ-TuRC, however, the presence of GIP-1 in the outer sphere associated with dynamic microtubules suggests that in *C. elegans* the γ-TuRC can assemble and function in the absence of MZT-1 as has been seen at other cellular sites (Sallee et al., 2018).

Our data suggest that PCM is initially removed from the centrosome by dephosphorylation, either through the direct action of PP2A phosphatases on PCM proteins or indirectly through the inactivation of mitotic kinases. This removal of PCM proteins from the inner sphere weakens the remaining PCM, allowing for rupture of the outer sphere by cortical pulling forces that rupture the remaining PCM into packets. The removal of both SPD-2 and MZT-1 appears to exclusively depend on phosphatase activity as they do not localize in packets and their disassembly was not affected by the inhibition of cortical forces.

Furthermore, a pool of both SPD-2 and SPD-5 remained at the centrosome following LET-92 depletion, indicating that the cortical forces alone are not sufficient for their effective clearance. Thus, PCM disassembly appears to be initiated by dephosphorylation by the PP2A subunit LET-92. As LET-92 plays a number of roles at the centrosome and phosphatase activity can directly regulate mitotic kinases (Enos et al., 2018; Kitagawa et al., 2011; Song et al., 2011), further studies will be necessary to determine if its role in PCM dissolution is direct or indirect.

Following dissolution, we found that the PCM fragments into small packets that retain MTOC potential. These packets are reminiscent of PCM flares described in Drosophila (Megraw, Kilaru, Turner, &Kaufman, 2002). Although PCM flares are reported to be present throughout the cell cycle rather than exclusively during centrosome disassembly as for the packets we describe, the molecular and mechanistic underpinnings of both of these structures might be common. For example, flares were first defined by their association with Centrosomin (Cnn), the proposed functional ortholog of SPD-5 (Megraw et al., 2002). Cnn is proposed to live in different states in the PCM in Drosophila, assembling first near the centrioles in a phosphorylated state and transiting towards the PCM periphery as a higher order multimerized scaffold where Cnn molecules are likely eventually dephosphorylated and lose PCM association (Conduit, Richens, et al., 2014b). Similarly, the inner sphere of SPD-5 may represent a specific pool of SPD-5 that can be readily dissociated by dephosphorylation, while the outer sphere may represent a macromolecular scaffold that relies on physical disruption for disassembly. Moreover, it appears from our observations that packets persist for several minutes in the cytoplasm before their complete disappearance, indicating a relatively stable state. Recent studies of *in vitro* assembled PCM point to different physical properties between ‘young’ and ‘old’ condensates of SPD-5, with young condensates behaving more like a liquid and old condensates acting more like a gel (Woodruff et al., 2017). Perhaps packets are the remnants of older gel-like matrices of SPD-5, which would also explain their ability to be torn apart by cortical forces.

Our results indicate that cortical forces can shape the PCM in multiple ways, mainly through an effect on outer sphere proteins. The balance of cortical forces appears to tune the levels of SPD-5 incorporation into the PCM, independently of SPD-2; decreasing or increasing cortical forces caused more or less SPD-5 incorporation but had no effect on the levels of SPD-2. Thus, cortical forces negatively regulate the growth of the PCM, hypothetically by physically removing PCM from the outer sphere. We found a pool of SPD-5 that remained at the centrosome after perturbation of cortical forces, further suggesting that SPD-5 can be differentially regulated within the PCM, perhaps through spatially segregated pools of differentially phosphorylated SPD-5.

In total, these results suggest that PCM is disassembled through the removal of the inner sphere of PCM by PP2A phosphatase activity, followed by the outer sphere by cortical pulling forces, which liberate dynamic microtubules and inactivate MTOC function at the centrosome. With an understanding of the mechanisms underlying this process, future studies will reveal whether hyperactive MTOC function at the centrosome has a direct effect on the cell cycle or cell differentiation in a developing organism, as has been previously postulated.

## Materials and Methods

### *C. elegans* strains and maintenance

*C. elegans* strains were maintained at 20°C unless otherwise specified and cultured as previously described (Brenner, 1974). Experiments were performed using embryos from one-day adults. Unless otherwise indicated, at least five embryos were scored in each experimental condition. Strains used in this study are as follows:

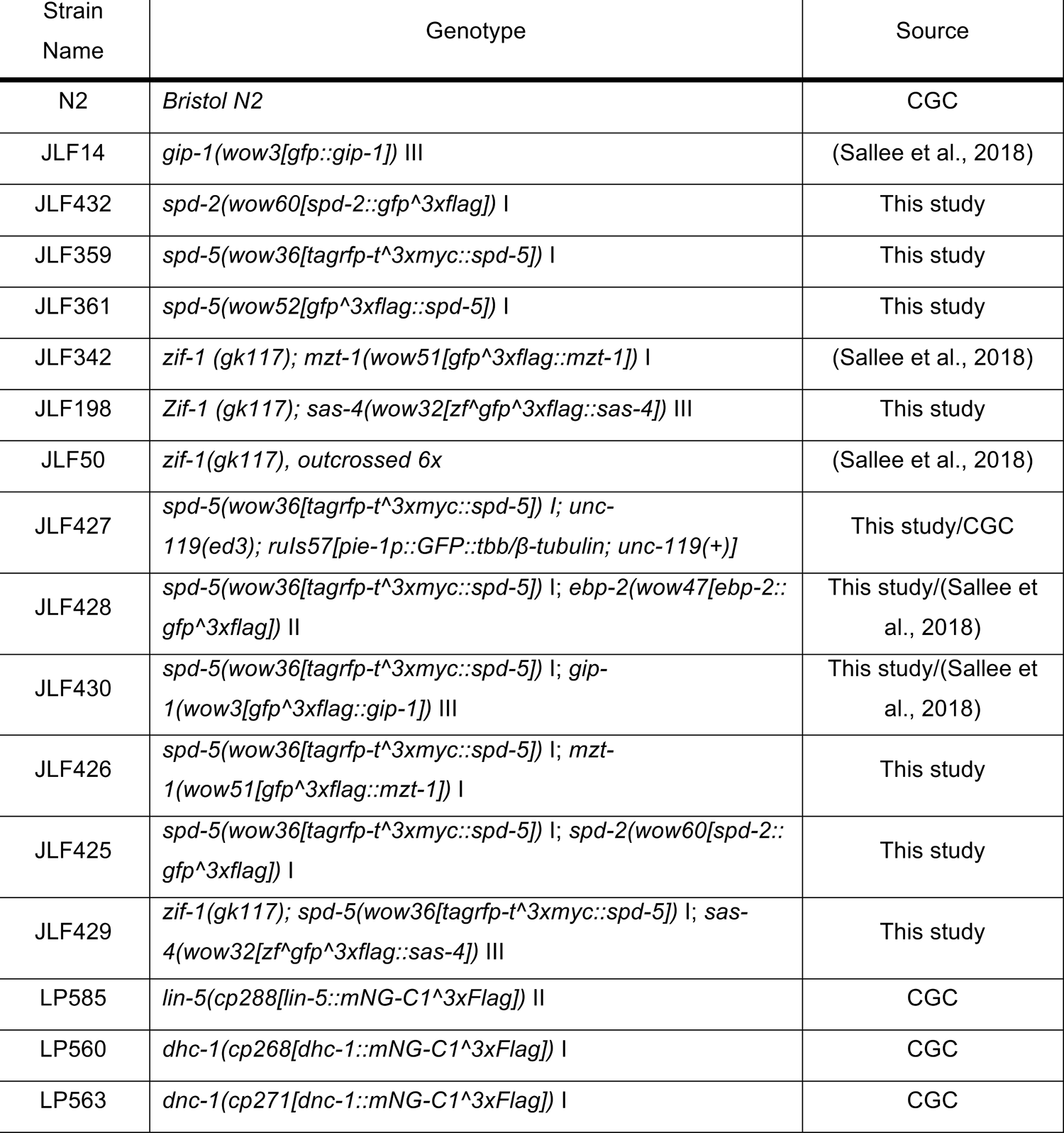

### CRISPR/Cas9

Endogenously tagged proteins used in this study were generated using the CRISPR Self Excising Cassette (SEC) method that has been previously described (Dickinson et al., 2015). DNA mixtures (sgRNA and Cas9 containing plasmid and repair template) were injected into young adults, and CRISPR edited worms were selected by treatment with hygromycin followed by visual inspection for appropriate expression and localization (Dickinson et al., 2015). sgRNA and homology arm sequences used to generate lines are as follows:

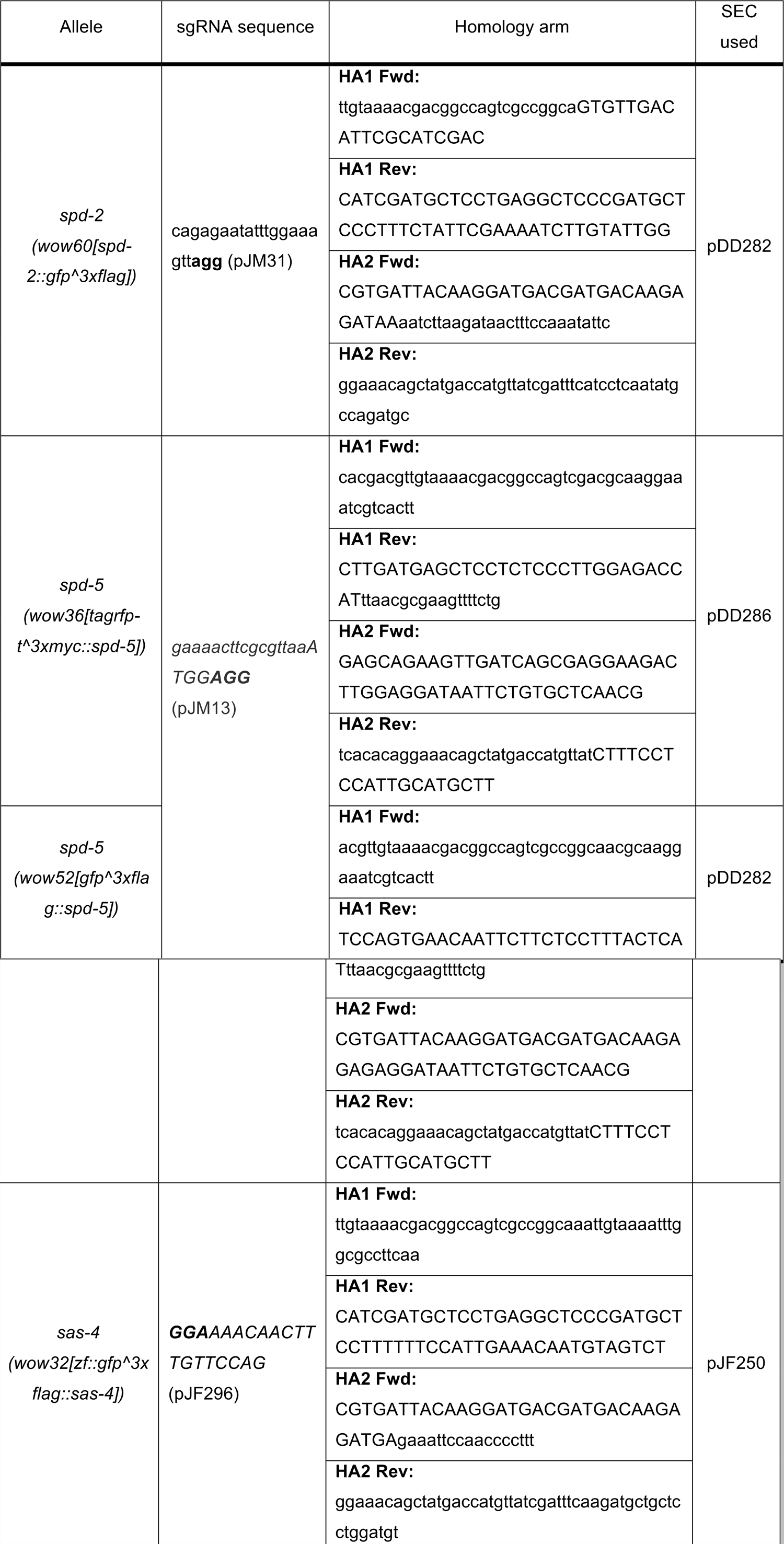

### Image acquisition

Embryos dissected from one-day old adults were mounted on a pad (3% agarose dissolved in M9) sandwiched between a microscope slide and no. 1.5 coverslip. Time-lapse images were acquired on a Nikon Ti-E inverted microscope (Nikon Instruments) equipped with a 1.5x magnifying lens, a Yokogawa X1 confocal spinning disk head, and an Andor Ixon Ultra back thinned EM-CCD camera (Andor), all controlled by NIS Elements software (Nikon). Images were obtained using a 60x Oil Plan Apochromat (NA=1.4) or 100x Oil Plan Apochromat (NA=1.45) objective. Z-stacks were acquired using a 0.5 μm step every minute. Images were adjusted for brightness and contrast using ImageJ software.

### Drug treatment

Drug treatments were performed as previously described (Yang &Feldman, 2015). Briefly, embryos were mounted between a slide and coverslip, supported with 22.5 μM beads (Whitehouse Scientific), and bathed in an osmotic control buffer (embryonic growth medium - EGM (Shelton &Bowerman, 1996)) supplemented with either 10% DMSO, 30 μM okadaic acid, or 60 μM rubratoxin A. Embryos were laser permeabilized at appropriate times using a Micropoint dye laser (coumarin 435nm) mounted on the spinning-disk confocal described above.

### RNAi treatment

RNAi treatment was performed by feeding as previously described using *csnk-1*(RNAi), *gpr-1/2*(RNAi), and *let-92*(RNAi) expressing HT115 bacteria from the Ahringer RNAi library (Ahringer, 2006; Fraser et al., 2000; Kamath et al., 2003). L4 stage worms were grown on RNAi plates (NGM supplemented with IPTG and Ampicillin) at 25°C for 24h-48h. RNAi plates were seeded with a bacterial culture grown overnight and subsequently grown 48h at room temperature protected from light.

### Image Quantification

#### PCM volume measurements

PCM volume was measured from stacks of images taken through the ABp centrosome closest to the coverslip at each timepoint. Image stacks were first processed to eliminate the cytosolic background by subtracting the mean intensity of 10 random points in the cytoplasm at each plane and each timepoint. Image stacks were then thresholded using the Otsu method (ImageJ) to delimit the PCM structure. Volume measurements were performed using the 3D object counter imageJ plugin (Bolte &Cordelières, 2006). Only the volume measured at the centrosome/centrioles was considered.

#### Intensity measurement

Total intensity was measured by defining an image stack 15 μm wide × 7.5 μm deep around the centrosome for each timepoint. Another stack of the exact same dimensions was generated in the cytoplasm. Both stacks were sum projected and the total intensity was measured by subtracting the total intensity of the cytoplasmic sum projection from the total intensity of the centrosome sum projection. Centrosomal intensity was calculated in the same way, but the ROI was selected manually following initial thresholding. Packet intensity was determined by removing a manually selected ROI for the centriole/centrosome. In Figure 5G, we accounted for the fact that *let-92* depletion results in centriole duplication defects in the one cell embryo (Song et al., 2011). In control embryos, we determined the average intensity of each of the two individual centriolar/centrosomal foci of either SPD-2 or SPD-5 at the end of disassembly (t = ∼5’). We compared this value to the average intensity of the single centrosomes in *let-92* depleted embryos at the end of disassembly (t = ∼15’). This type of measurement was in contrast to the total centriole/centrosome measurement shown in Figure 5E and F, which does not distinguish the two resulting centrioles/centrosomes in control conditions at the end of disassembly.

#### Timing of events

The different steps of disassembly were defined based on hallmarks of both volume and intensity measurements. ‘Dissolution’ was defined as the timepoint at which the first decrease in PCM intensity was detected, which corresponded to a decrease in SPD-2 intensity. ‘Rupture’ was defined as the timepoint at which the first decrease in PCM volume was detected, which corresponded to a drop in SPD-5 volume. Packet formation was defined as the timepoint at which individualized foci of SPD-5 appeared around the centrioles.

#### Statistics

Statistical analyses were performed using R and Prism (GraphPad software, La Jolla, Ca, USA).

## Acknowledgements

We thank Kevin O’Connell, Jyoti Iyer, Dan Dickinson, and Bob Goldstein for CRISPR advice and protocols. We also thank Tim Stearns, Ariana Sanchez, Maria Sallee, Melissa Pickett and members of the Feldman lab for helpful discussions about the manuscript. Some of the nematode strains used in this work were provided by the *Caenorhabditis* Genetic Center, which is funded by the NIH Office of Research Infrastructure Programs (P40 OD010440). This work was supported by a March of Dimes Basil O’Connor Starter Scholar Research Award and an NIH New Innovator Award DP2GM119136-01 awarded to J.L.F. J.M. is supported by an American Heart Postdoctoral Fellowship.

## Author Contributions

Conceptualization, J.L.F and J.M.; Methodology, J.M., J.L.F., and J.C.Z.; Formal Analysis, J.M.; Investigation, J.M., J.L.F., and J.C.Z.; Writing - Original Draft, J.L.F and J.M.; Writing - Review &Editing, J.L.F and J.M.; Visualization, J.L.F and J.M.; Supervision, J.L.F.; Funding Acquisition, J.L.F and J.M.

## Supplemental Information

### Supplemental Figure Legends

**Figure 1-figure supplement 1.**
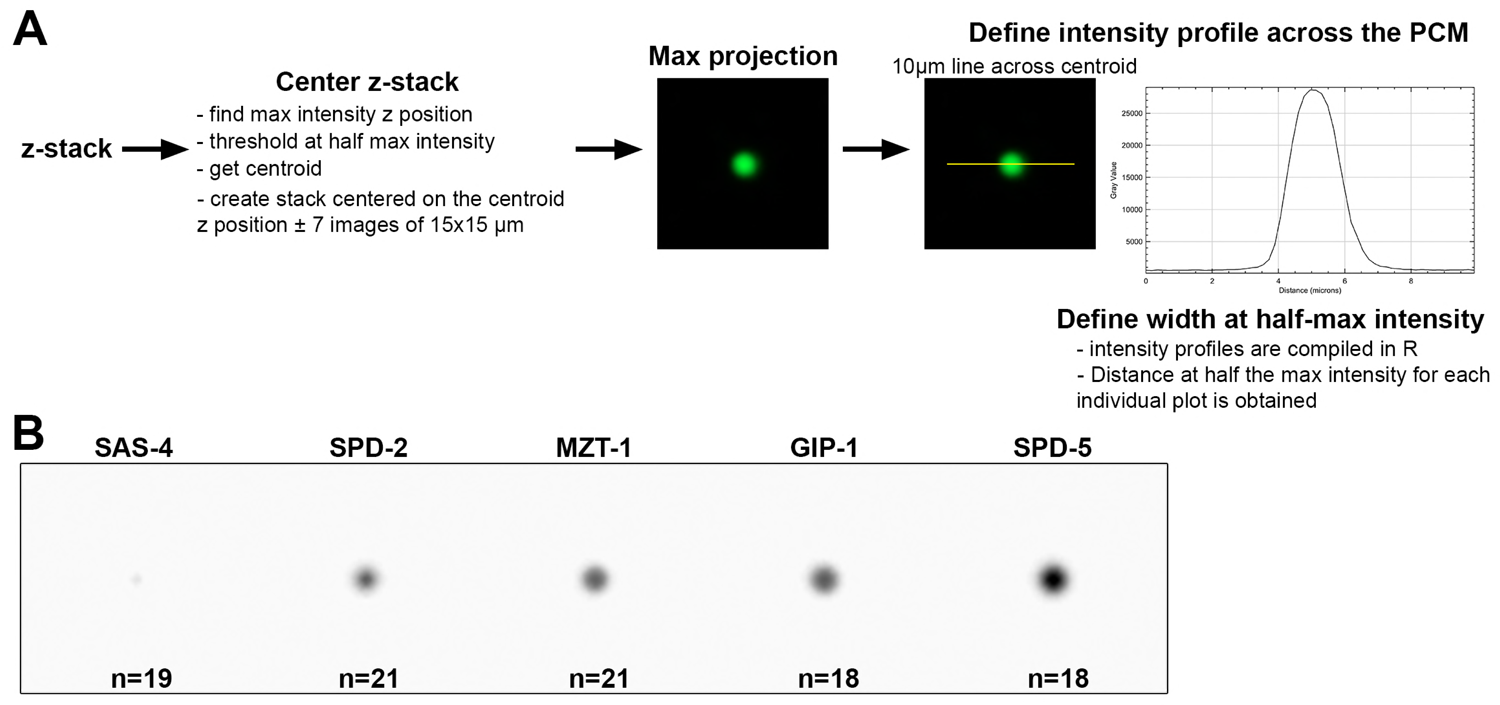
Methods for quantifying PCM width. A) The width of each PCM protein was determined using the same image analysis pipeline. Image stacks of about 30 images separated by 0.5 μm z-steps (15μm total) were acquired at NEBD using the same imaging parameters. Stacks were then cropped to include only the ABp cell and centered around the ABp centrosome closest to the coverslip. We found the max intensity ABp centrosome slice and created a new 30μm wide substack centered around this slice ± 7 slices (15 slices, 7μm total). The max intensity slice was then used to find the centroid of the centrosomal structure. This slice was thresholded using the half max intensity and the centroid value was obtained using the Analyze Particle tool (ImageJ). Using the coordinates of the centroid (X,Y) and the max intensity (Z), we created a 15μm wide substack centered on those coordinates ± 7 slices. The intensity profile was obtained by drawing a 10μm long line centered on the centroid. Profiles for each embryo were compiled and for each of them the width was determined by measuring the distance at the half max intensity. B) Mean projection of all max projections used in the study for measuring protein width.

**Figure 4-figure supplement 1.**
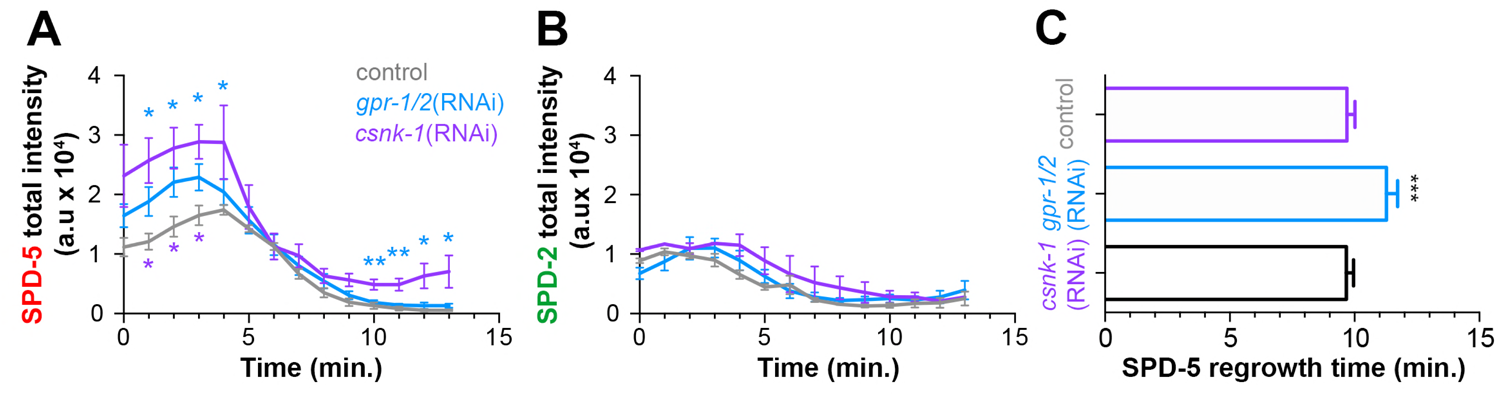
Cortical forces affect SPD-5, but not SPD-2, intensity and regrowth in the next cell cycle. Total SPD-5 (A) or SPD-2 (B) intensity at the centrosome during disassembly starting at NEBD (t=0 min) in embryos treated with lacZ(RNAi) (control, grey), *gpr-1/2(RNAi)* (blue), or *csnk-1(RNAi)* (purple). *p-value, gpr-1/2(RNAi) or *csnk-* 1(RNAi) vs. control < 0.05. ** p-value, gpr-1/2(RNAi) or csnk-1(RNAi) vs. control < 0.01. C) Average time after NEBD before regrowth of SPD-5 at the centrosome in the next cell cycle. Control: 9.69±0.33, n=8; *gpr-1/2(RNAi):* 11. 27±0.45, n=11; *csnk-1(RNAi):* 9.67±0.29, n=7. ***p-value, control or *csnk-1(RNAi)* vs. *gpr-1/2(RNAi)*, < 0.0001. ‘a.u.’ = arbitrary units. Error bars indicate standard deviation of the mean.

**Figure 4-figure supplement 2.**
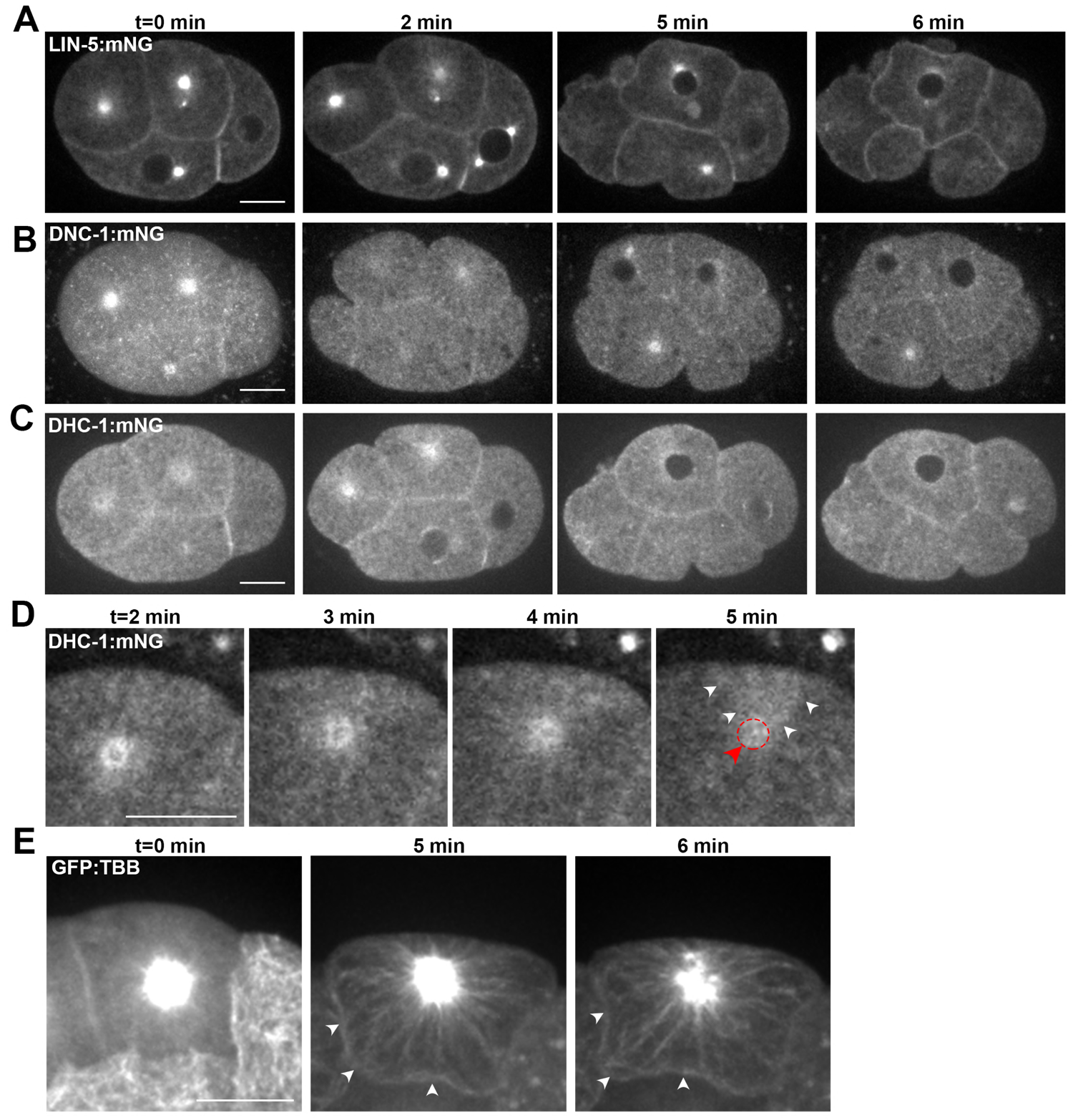
Localization of cortical force generating proteins and astral microtubules during PCM disassembly. A-C) Time lapse analysis of single plane images from 4-cell embryos expressing endogenous LIN-5::mNG (A), DNC-1::mNG (B) and DHC-1::mNG (C). D) Single plane image of the ABp cell division starting at NEBD (t=0 min) in an embryo expressing endogenous DHC-1::GFP. Movement of DHC-1 from the centrosome (red arrow and dotted circle) toward the dorsal membrane (white arrowheads) is apparent. E) Time lapse analysis of embryo expressing pie-1::TBB-2/ß>-tubulin::GFP starting at NEBD (t= 0 min, left panel) and 5 min (middle panel) and 6 min (right panel) after. Scale bars, 10μm.

### Video Legends

**Video 1.** Centrosome disassembly in the ABp cell in a 4-cell embryo expressing endogenous SPD-2::GFP. Scale bar, 5μm.

**Video 2.** Centrosome disassembly in the ABp cell in a 4-cell embryo expressing endogenous GFP::MZT-1. Scale bar, 5μm.

**Video 3.** Centrosome disassembly in the ABp cell in a 4-cell embryo expressing endogenous GFP::SPD-5. Yellow arrowhead and ‘c’ mark the centrioles. White arrowhead and ‘p’ mark the packets. Scale bar, 5μm.

**Video 4.** Centrosome disassembly in the ABp cell in a 4-cell embryo expressing endogenous GFP::GIP-1. Yellow arrowhead and ‘c’ mark the centrioles. White arrowhead and ‘p’ mark the packets. Scale bars, 5μm.

**Video 5.** Centrosome disassembly in the ABp cell in a 4-cell embryo expressing endogenous tagRFP-T::SPD-5; GFP::GIP-1. Yellow arrowhead and ‘c’ mark the centrioles. White arrowhead and ‘p’ mark the packets. Scale bars, 5μm.

## References

Ahringer, J. (2006). Reverse genetics. WormBook. http://doi.Org/10.1895/wormbook.1.47.1

Bobinnec, Y., Fukuda, M., & Nishida, E. (2000). Identification and characterization of Caenorhabditis elegans gamma-tubulin in dividing cells and differentiated tissues. Journal of Cell Science, 113 Pt 21(DECEMBER 2000), 3747–3759.

Bolte, S., & Cordelières, F. P. (2006). A guided tour into subcellular colocalization analysis in light microscopy. Journal of Microscopy, 224(Pt 3), 213–232. http://doi.org/10.1111/j.1365-2818.2006.01706.x

Borrego-Pinto, J., Somogyi, K., Karreman, M. A., König, J., Müller-Reichert, T., Bettencourt-Dias, M., et al. (2016). Distinct mechanisms eliminate mother and daughter centrioles in meiosis of starfish oocytes. The Journal of Cell Biology, 212(7), 815–827. http://doi.org/10.1083/jcb.201510083

Brenner, S. (1974). The genetics of Caenorhabditis elegans. Genetics, 77(1), 71–94.

Cabral, G., Sans, S. S., Cowan, C. R., & Dammermann, A. (2013). Multiple mechanisms contribute to centriole separation in C. elegans. Current Biology: CB, 23(14), 1380–1387. http://doi.org/10.1016/j.cub.2013.06.043

Cheng, J., Tiyaboonchai, A., Yamashita, Y. M., & Hunt, A. J. (2011). Asymmetric division of cyst stem cells in Drosophila testis is ensured by anaphase spindle repositioning. Development (Cambridge, England), 138(5), 831–837. http://doi.org/10.1242/dev.057901

Chichinadze, K., Lazarashvili, A., & Tkemaladze, J. (2012). RNA in centrosomes: Structure and possible functions. Protoplasma, 250(1), 397–405. http://doi.org/10.1007/s00709-012-0422-6

Conduit, P. T., & Raff, J. W. (2010). Cnn dynamics drive centrosome size asymmetry to ensure daughter centriole retention in Drosophila neuroblasts. Current Biology: CB, 20(24), 2187–2192. http://doi.org/10.1016/j.cub.2010.11.055

Conduit, P. T., Brunk, K., Dobbelaere, J., Dix, C. I., Lucas, E. P., & Raff, J. W. (2010). Centrioles regulate centrosome size by controlling the rate of Cnn incorporation into the PCM. Current Biology: CB, 20(24), 2178–2186. http://doi.org/10.1016Zj.cub.2010.11.011

Conduit, P. T., Feng, Z., Richens, J. H., Baumbach, J., Wainman, A., Bakshi, S. D., et al. (2014a). The centrosome-specific phosphorylation of Cnn by Polo/Plk1 drives Cnn scaffold assembly and centrosome maturation. Developmental Cell, 28(6), 659–669. http://doi.org/10.1016/j.devcel.2014.02.013

Conduit, P. T., Richens, J. H., Wainman, A., Holder, J., Vicente, C. C., Pratt, M. B., et al. (2014b). A molecular mechanism of mitotic centrosome assembly in Drosophila. eLife, 3, e03399. http://doi.org/10.7554/eLife.03399

Decker, M., Jaensch, S., Pozniakovsky, A., Zinke, A.,O’Connell, K. F., Zachariae, W., et al. (2011). Limiting amounts of centrosome material set centrosome size in C. elegans embryos. Current Biology: CB, 21(15), 1259–1267. http://doi.org/10.1016Zj.cub.2011.06.002

Dickinson, D. J., Pani, A. M., Heppert, J. K., Higgins, C. D., & Goldstein, B. (2015). Streamlined Genome Engineering with a Self-Excising Drug Selection Cassette. Genetics, 200(4), 1035–1049. http://doi.org/10.1534/genetics.115.178335

Dickinson, D. J., Ward, J. D., Reiner, D. J., & Goldstein, B. (2013). Engineering the Caenorhabditis elegans genome using Cas9-triggered homologous recombination. Nature Methods, 10(10), 1028–1034. http://doi.org/10.1038/nmeth.2641

Dictenberg, J. B., Zimmerman, W., Sparks, C. A., Young, A., Vidair, C., Zheng, Y., et al. (1998). Pericentrin and gamma-tubulin form a protein complex and are organized into a novel lattice at the centrosome. The Journal of Cell Biology, 141(1), 163–174.

Enos, S. J., Dressler, M., Gomes, B. F., Hyman, A. A., & Woodruff, J. B. (2018). Phosphatase PP2A and microtubule-mediated pulling forces disassemble centrosomes during mitotic exit. Biology Open, 7(1), bio029777. http://doi.org/10.1242/bio.029777

Fraser, A. G., Kamath, R. S., Zipperlen, P., Martinez-Campos, M., Sohrmann, M., & Ahringer, J. (2000). Functional genomic analysis of C. elegans chromosome I by systematic RNA interference. Nature, 408(6810), 325–330. http://doi.org/10.1038/35042517

Fry, A. M., Sampson, J., Shak, C., & Shackleton, S. (2017). Recent advances in pericentriolar material organization: ordered layers and scaffolding gels. F1000Research, 6, 1622–10. http://doi.org/10.12688/f1000research.11652.1

Fu, J., & Glover, D. M. (2012). Structured illumination of the interface between centriole and peri-centriolar material. Open Biology, 2(8), 120104–120104. http://doi.org/10.1098/rsob.120104

Godinho, S. A., & Pellman, D. (2014). Causes and consequences of centrosome abnormalities in cancer, 369(1650), 20130467–20130467. http://doi.org/10.1098/rstb.2013.0467

Gonczy, P., & Rose, L. S. (2005). Asymmetric cell division and axis formation in the embryo. WormBook, 1–20. http://doi.org/10.1895/wormbook.130.1

Grill, S. W., Gonczy, P., Stelzer, E. H., & Hyman, A. A. (2001). Polarity controls forces governing asymmetric spindle positioning in the Caenorhabditis elegans embryo. Nature, 409(6820), 630–633. http://doi.org/10.1038/35054572

Hamill, D. R., Severson, A. F., Carter, J. C., & Bowerman, B. (2002). Centrosome maturation and mitotic spindle assembly in C. elegans require SPD-5, a protein with multiple coiled-coil domains. Developmental Cell, 3(5), 673–684.

Hannak, E., Oegema, K., Kirkham, M., Gonczy, P., Habermann, B., & Hyman, A. A. (2002). The kinetically dominant assembly pathway for centrosomal asters in Caenorhabditis elegans is gamma-tubulin dependent. The Journal of Cell Biology, 157(4), 591–602. http://doi.org/10.1083/jcb.200202047

Kamath, R. S., Fraser, A. G., Dong, Y., Poulin, G., Durbin, R., Gotta, M., et al. (2003). Systematic functional analysis of the Caenorhabditis elegans genome using RNAi. Nature, 421(6920), 231–237. http://doi.org/10.1038/nature01278

Kemp, C. A., Kopish, K. R., Zipperlen, P., Ahringer, J., & O’Connell, K. F. (2004). Centrosome maturation and duplication in C. elegans require the coiled-coil protein SPD-2. Developmental Cell, 6(4), 511–523. http://doi.org/10.1016/S1534-5807(04)00066-8

Kitagawa, D., Flückiger, I., Polanowska, J., Keller, D., Reboul, J., & Gönczy, P. (2011). PP2A phosphatase acts upon SAS-5 to ensure centriole formation in C. elegans embryos. Developmental Cell, 20(4), 550–562. http://doi.org/10.1016Zj.devcel.2011.02.005

Kotak, S., & Gönczy, P. (2013). Mechanisms of spindle positioning: cortical force generators in the limelight. Current Opinion in Cell Biology, 25(6), 741–748. http://doi.org/10.1016/j.ceb.2013.07.008

Lawo, S., Hasegan, M., Gupta, G. D., & Pelletier, L. (2012). Subdiffraction imaging of centrosomes reveals higher-order organizational features of pericentriolar material. Nature Cell Biology, 14(11), 1148–1158. http://doi.org/10.1038/ncb2591

Lin, T.-C., Neuner, A., & Schiebel, E. (2015). Targeting of **γ**-tubulin complexes to microtubule organizing centers: conservation and divergence. Trends in Cell Biology, 25(5), 296–307. http://doi.org/10.1016/j.tcb.2014.12.002

Lingle, W. L., Lutz, W. H., Ingle, J. N., Maihle, N. J., & Salisbury, J. L. (1998). Centrosome hypertrophy in human breast tumors: implications for genomic stability and cell polarity. Proceedings of the National Academy of Sciences, 95(6), 2950–2955.

Lu, Y., & Roy, R. (2014). Centrosome/Cell cycle uncoupling and elimination in the endoreduplicating intestinal cells of C. elegans. PLoS ONE, 9(10), e110958. http://doi.org/10.1371/journal.pone.0110958

Luksza, M., Queguigner, I., Verlhac, M.-H., & Brunet, S. (2013). Rebuilding MTOCs upon centriole loss during mouse oogenesis. Developmental Biology, 382(1), 48–56. http://doi.org/10.1016/j.ydbio.2013.07.029

Megraw, T. L., Kilaru, S., Turner, F. R., & Kaufman, T. C. (2002). The centrosome is a dynamic structure that ejects PCM flares. Journal of Cell Science, 115(Pt 23), 4707–4718.

Mennella, V., Agard, D. A., Huang, B., & Pelletier, L. (2014). Amorphous no more: subdiffraction view of the pericentriolar material architecture. Trends in Cell Biology, 24(3), 188–197. http://doi.org/10.1016/j.tcb.2013.10.001

Mennella, V., Keszthelyi, B., McDonald, K. L., Chhun, B., Kan, F., Rogers, G. C., et al. (2012). Subdiffraction-resolution fluorescence microscopy reveals a domain of the centrosome critical for pericentriolar material organization. Nature Cell Biology, 14(11), 1159–1168. http://doi.org/10.1038/ncb2597

Mikeladze-Dvali, T., Tobel, von, L., Strnad, P., Knott, G., Leonhardt, H., Schermelleh, L., & Gonczy, P. (2012). Analysis of centriole elimination during C. elegans oogenesis, 139(9), 1670–1679. http://doi.org/10.1242/dev.075440

Motegi, F., Velarde, N. V., Piano, F., & Sugimoto, A. (2006). Two phases of astral microtubule activity during cytokinesis in C. elegans embryos. Developmental Cell, 10(4), 509–520. http://doi.org/10.1016Zj.devcel.2006.03.001

Muroyama, A., Seldin, L., & Lechler, T. (2016). Divergent regulation of functionally distinct y-tubulin complexes during differentiation. The Journal of Cell Biology, 213(6), 679–692. http://doi.org/10.1083/jcb.201601099

Novak, Z. A., Wainman, A., Gartenmann, L., & Raff, J. W. (2016). Cdk1 Phosphorylates Drosophila Sas-4 to Recruit Polo to Daughter Centrioles and Convert Them to Centrosomes. Developmental Cell, 37(6), 545–557. http://doi.org/10.1016/j.devcel.2016.05.022

Oakley, B. R., Paolillo, V., & Zheng, Y. (2015). γ-Tubulin complexes in microtubule nucleation and beyond. Molecular Biology of the Cell, 26(17), 2957–2962. http://doi.org/10.1091/mbc.E14-11-1514

Pihan, G. A. (2013). Centrosome dysfunction contributes to chromosome instability, chromoanagenesis, and genome reprograming in cancer. Frontiers in Oncology, 3, 277. http://doi.org/10.3389/fonc.2013.00277

Pihan, G. A., Purohit, A., Wallace, J., Malhotra, R., Liotta, L., & Doxsey, S. J. (2001). Centrosome defects can account for cellular and genetic changes that characterize prostate cancer progression. Cancer Research, 61(5), 2212–2219.

Pimenta-Marques, A., Bento, I., Lopes, C. A. M., Duarte, P., Jana, S. C., & Bettencourt-Dias, M. (2016). A mechanism for the elimination of the female gamete centrosome in Drosophila melanogaster. Science, 353(6294), aaf4866–aaf4866. http://doi.org/10.1126/science.aaf4866

Rose, L., & Gönczy, P. (2014). Polarity establishment, asymmetric division and segregation of fate determinants in early C. elegans embryos. WormBook, 1–43. http://doi.org/10.1895/wormbook.1.30.2

Salisbury, J. L., Lingle, W. L., White, R. A., Cordes, L. E., & Barrett, S. (1999). Microtubule nucleating capacity of centrosomes in tissue sections. The Journal of Histochemistry and Cytochemistry: Official Journal of the Histochemistry Society, 47(10), 1265–1274. http://doi.org/10.1177/002215549904701006

Sallee, M. D., Zonka, J. C., Skokan, T. D., Raftrey, B. C., & Feldman, J. L. (2018). Tissue-specific degradation of essential centrosome components reveals distinct microtubule populations at microtubule organizing centers. PLoS Biology, 16(8), e2005189-32. http://doi.org/10.1371/journal.pbio.2005189

Sanchez, A. D., & Feldman, J. L. (2016). Microtubule-organizing centers: from the centrosome to non-centrosomal sites. Current Opinion in Cell Biology. http://doi.org/10.1016/j.ceb.2016.09.003

Schlaitz, A.-L., Srayko, M., Dammermann, A., Quintin, S., Wielsch, N., MacLeod, I., et al. (2007). The C. elegans RSA complex localizes protein phosphatase 2A to centrosomes and regulates mitotic spindle assembly. Cell, 128(1), 115–127. http://doi.org/10.1016Zj.cell.2006.10.050

Shelton, C. A., & Bowerman, B. (1996). Time-dependent responses to glp-1-mediated inductions in early C. elegans embryos. Development, 122(7), 2043–2050.

Song, M. H., Liu, Y., Anderson, D. E., Jahng, W. J., & O'Connell, K. F. (2011). Protein phosphatase 2A-SUR-6/B55 regulates centriole duplication in C. elegans by controlling the levels of centriole assembly factors. Developmental Cell, 20(4), 563–571. http://doi.org/10.1016/j.devcel.2011.03.007

Stubenvoll, M. D., Medley, J. C., Irwin, M., & Song, M. H. (2016). ATX-2, the C. elegans Ortholog of Human Ataxin-2, Regulates Centrosome Size and Microtubule Dynamics. PLoS Genetics, 12(9), e1006370. http://doi.org/10.1371/journal.pgen.1006370

Tsou, M.-F. B., & Stearns, T. (2006). Mechanism limiting centrosome duplication to once per cell cycle. Nature, 442(7105), 947–951. http://doi.org/10.1038/nature04985

Woodruff, J. B., Gomes, B. F., Widlund, P. O., Mahamid, J., Honigmann, A., & Hyman, A. A. (2017). The Centrosome Is a Selective Condensate that Nucleates Microtubules by Concentrating Tubulin. Cell, 169(6), 1066–1071.e10. http://doi.org/10.1016/j.cell.2017.05.028

Woodruff, J. B., Wueseke, O., & Hyman, A. A. (2014). Pericentriolar material structure and dynamics. Philosophical Transactions of the Royal Society of London. Series B, Biological Sciences, 369(1650). http://doi.org/10.1098/rstb.2013.0459

Woodruff, J. B., Wueseke, O., Viscardi, V., Mahamid, J., Ochoa, S. D., Bunkenborg, J., et al. (2015). Regulated assembly of a supramolecular centrosome scaffold in vitro. Science, 348(6236), 808–812. http://doi.org/10.1126/science.aaa3923

Wueseke, O., Bunkenborg, J., Hein, M. Y., Zinke, A., Viscardi, V., Woodruff, J. B., Oegema, K., Mann, M., Andersen, J. S., & Hyman, A. A. (2014a). The Caenorhabditis elegans pericentriolar material components SPD-2 and SPD-5 are monomeric in the cytoplasm before incorporation into the PCM matrix. Molecular Biology of the Cell, 25(19), 2984–2992. http://doi.org/10.1091/mbc.E13-09-0514

Wueseke, O., Bunkenborg, J., Hein, M. Y., Zinke, A., Viscardi, V., Woodruff, J. B., Oegema, K., Mann, M., Andersen, J. S., & Hyman, A. A. (2014b). The Caenorhabditis elegans pericentriolar material components SPD-2 and SPD-5 are monomeric in the cytoplasm before incorporation into the PCM matrix. Molecular Biology of the Cell, 25(19), 2984–2992. http://doi.org/10.1091/mbc.E13-09-0514

Wueseke, O., Zwicker, D., Schwager, A., Wong, Y. L., Oegema, K., Jülicher, F., et al. (2016). Polo-like kinase phosphorylation determines Caenorhabditis elegans centrosome size and density by biasing SPD-5 toward an assembly-competent conformation. Biology Open, 5(10), 1431–1440. http://doi.org/10.1242/bio.020990

Yang, R., & Feldman, J. L. (2015). SPD-2/CEP192 and CDK Are Limiting for Microtubule-Organizing Center Function at the Centrosome. Current Biology: CB, 25(14), 1924–1931. http://doi.org/10.1016/j.cub.2015.06.001

Zebrowski, D. C., Vergarajauregui, S., Wu, C.-C., Piatkowski, T., Becker, R., Leone, M., et al. (2015). Developmental alterations in centrosome integrity contribute to the post-mitotic state of mammalian cardiomyocytes. eLife, 4, 461. http://doi.org/10.7554/eLife.05563

